# Competitive release in tumors

**DOI:** 10.1101/263335

**Authors:** Yongqian Ma, Jeffrey West, Paul K. Newton

## Abstract

Competitive release is a bedrock principle of coevolutionary ecology and population dynamics. It is also the main mechanism by which heterogeneous tumors develop chemotherapeutic resistance. Understanding, controlling, and exploiting this important mechanism represents one of the key challenges and potential opportunities of current medical oncology. The development of sophisticated mathematical and computational models of coevolution among clonal and sub-clonal cell populations in the tumor ecosystem can guide us in predicting and shaping various responses to perturbations in the fitness landscape which is altered by chemo-toxic agents. This in turn can help us design adaptive chemotherapeutic strategies to combat the release resistant cells.

## I. INTRODUCTION TO THE TYPE OF PROBLEM IN CANCER

The important mechanism of *competitive release* has its roots in ecology [1] and population dynamics [2] where sub-populations compete for resources as they coevolve in a resource limited microenvironment. Imagine one species (e.g. Connell’s blue barnacle species [1] normally occupying the intertidal zone) competing with another (the brown barnacle species normally occupying the coast region above high tide). The brown barnacles cannot colonize the intertidal zone because the blue barnacles outcompete them for resources in that ecological niche. By removing the blue barnacles, the browns can flourish because they are *released from the previous competition* which was keeping their population in check [1]. This important and well documented coevolutionary mechanism is used effectively for pest control in managing sub-populations of damaging insects (called integrated pest management). By keeping just enough of a more fit species present, other more invasive species can sometimes be successfully controlled with restricted population growth strategies without ever actually eliminating them. Effective use of this concept relies on both a working understanding of how the many interacting sub-species coevolve, as well as the ability to continually monitor the different sub-populations so that adaptive adjustments can be made to the delivery schedule of toxins to shape the fitness landscape on a timescale shorter than the timescale on which the population develops resistance to the pesticide.

Many of these same ideas can also be used for the control of the heterogeneous population of cancer cells comprising an evolving and growing tumor [2-10]. The concept, known generically as adaptive therapeutics [10] is an exciting avenue for avoiding chemo-resistance in a tumor which is one of the main stumbling blocks associated with the efficacy of many cancer treatments. If a tumor was made up of a homogeneous population of identical cells, a clear strategy would be to kill as many as possible using the maximum tolerated dose [11]. We know, however, that a typical tumor ecosystem is instead comprised of a heterogeneous coevolving population of sub-clones [5,7,9,12]. What are the strategic implications of this fact? For simplicity, consider a system of three coevolving populations of cells: healthy (H), chemo-sensitive (S) (proliferative), and chemo-resistant (R) (non-proliferative), as shown in Figure 1. A tumor initially grows because of uncontrolled division by the proliferative sub-population. Often, there is a smaller pre-existing sub-population of chemo-resistant cells in the tumor that are held in check by the more aggressive and rapidly expanding population of chemo-sensitive cells of higher fitness [2,5,13]. After one or more rounds of chemotherapy, the chemo-sensitive cell population is drastically reduced or possibly even eliminated, selecting for the chemo-resistant sub-population. The next rounds of chemotherapy will no longer be effective on these chemo-resistant cells and treatment will fail. Coevolutionary mathematical models using either replicator dynamics (relative proportion of cell sub-types) or Moran processes (finite populations) with fitness determined by payoff matrices associated with evolutionary games [13-15] form the basis for most of these models. We describe results using the Prisoner’s Dilemma evolutionary game [16] for cell-cell interactions where the healthy cells are the cooperators, and the cancer cells are the defectors [16-18]. More complex models have been developed that include spatial effects [19], effects of the tumor microenvironment [20], and immune responses [21]. Evolutionary models producing simulated clinical trials are increasingly being developed as a way of predicting future hypothetical tumor states to guide and inform personalized treatment strategies [22].

**Figure 1.**
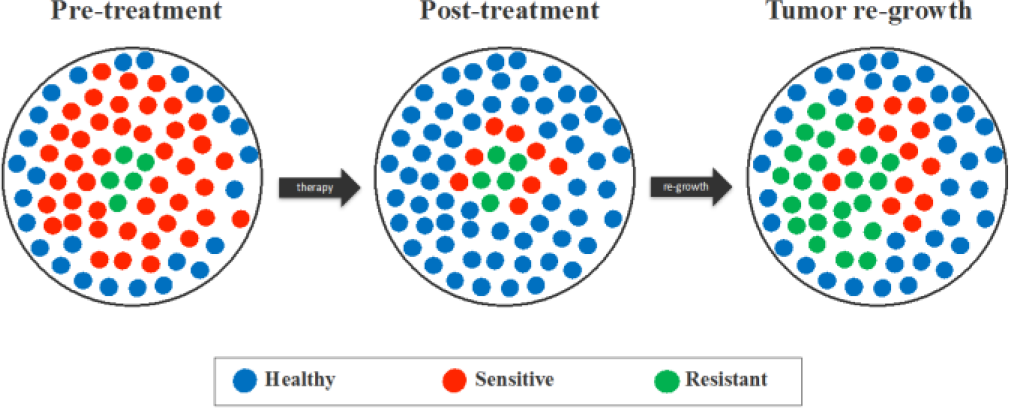
Competitive release in a tumor. Before treatment (left), the sensitive cells dominate the tumor, with a smaller pre-exisiting resistant sub-population. Chemotherapy kills off most of the sensitive cells (middle), thus selecting for the resistant strain. The relative fitness landscape has changed, and the chemo-resistant cells flourish (right), rendering the subsequent rounds of treatment ineffective.

## II. ILLUSTRATIVE RESULTS OF APPLICATION OF METHODS

Figure 2 shows how the fitness landscape is altered by the application of chemotherapy. With no therapy, the sensitive cells have a fitness advantage over the resistant and healthy cells, keeping them in check. Chemotherapy alters the fitness landscape, selecting for the resistant cell type (green) which gets released and subsequently dominates the tumor. A typical simulation is shown in Figure 3, using the Prisoner’s Dilemma (4) payoff matrix for cell-cell interactions. With a continuous dose of chemotherapy, the resistant cells are released and can repopulate the tumor, making future rounds of chemotherapy ineffective.

**Figure 2.**
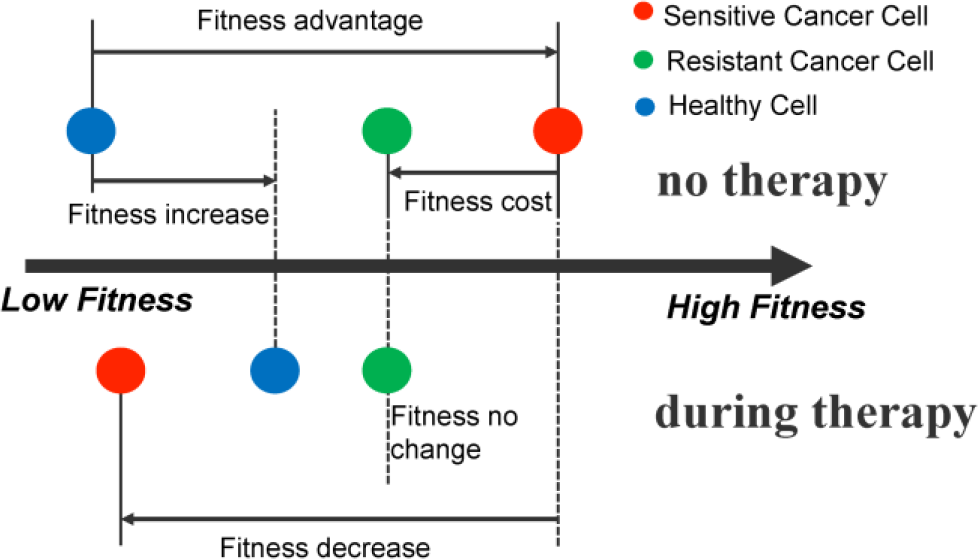
Phenotypic cost of resistance. A schematic of the fitness of each subpopulation before therapy (top) and during therapy (bottom). A driver mutation leads to a fitness advantage of the cancer cell (red), determined by the Prisoner′s Dilemma. A subsequent resistant-conferring mutation comes at a cost (green). The fitness of the resistant population is unaffected by therapy′s selective pressure, but the healthy population is given an advantage over the chemo-sensitive population.

**Figure 3.**
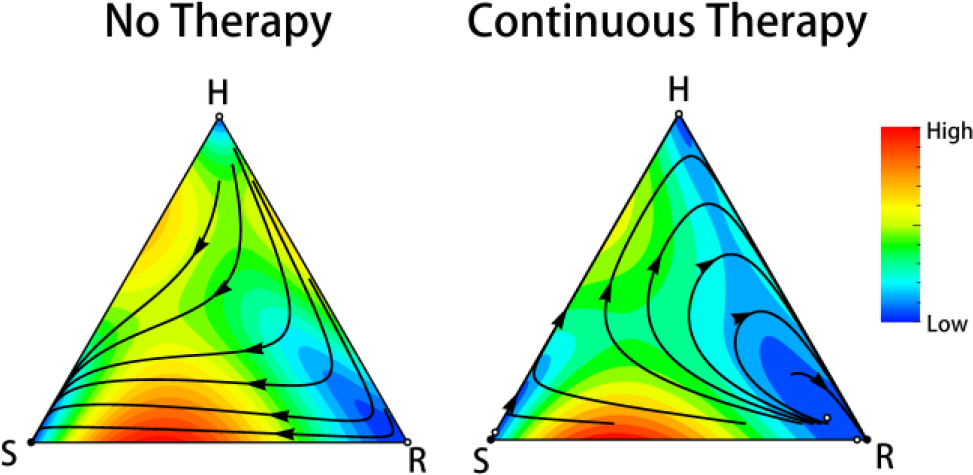
Competitive release in action. With no therapy (left), the sensitive cells have a fitness advantage over both the resistant cells and the healthy cells, so all initial distributions of cells grow into a tumor dominated by sensitive cells. With continuous therapy (right), the fitness of the sensitive cells is reduced, while the resistant cells have highest fitness and are released and allowed to grow unchallenged. At the end of the treatment cycle, the bulk tumor is made up of chemo-resistant cells.

## III. QUICK GUIDE TO THE METHODS

The dynamics are controlled by the replicator equations (with no mutations) [16]:

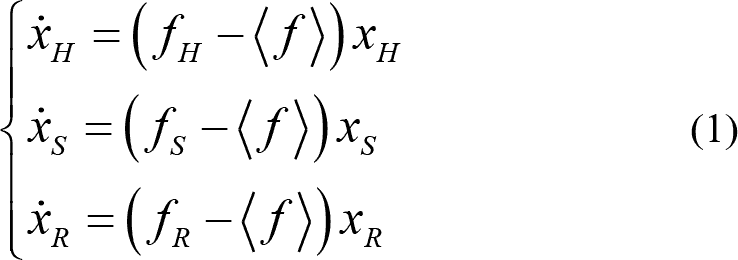

Fitness functions for Healthy (H), Sensitive (S), and Resistant (R) cells are given by:

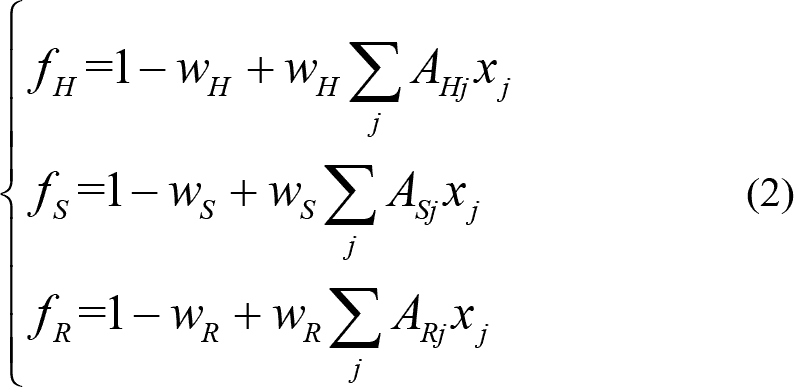

The average system fitness is:

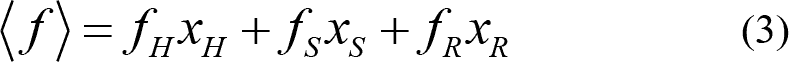

The parameters 0≤*w_H_*≤1;0≤*w_S_*≤1;0≤*w_R_*≤1 are used to specify the selection pressure on each of the cell types, and are adjusted as a way of simulating the effects of chemo-toxic agents [18].

The 3×3 payoff matrix *A* is of Prisoner’s dilemma type [13,16,17]:

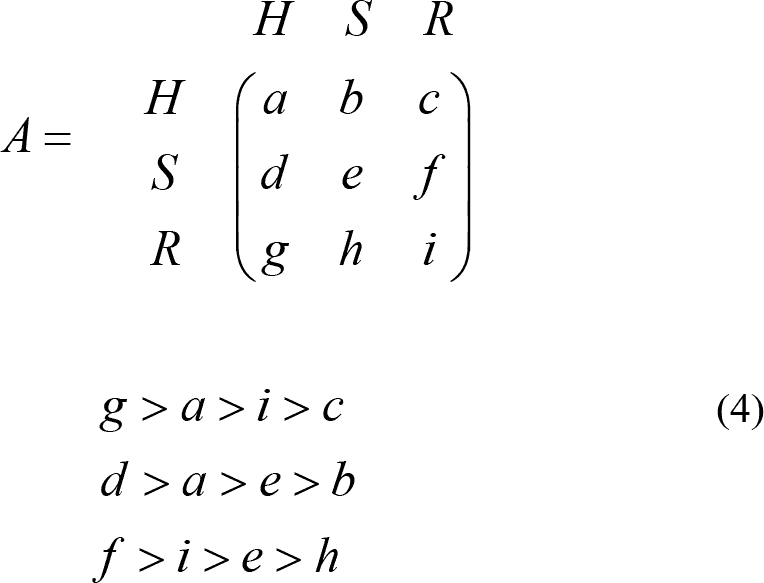

